# Does deliberation really need more effort than intuition? A test using event-related brain potential

**DOI:** 10.1101/628446

**Authors:** Jianmin Zeng, Dinglan Tang, Jia Yu, Qinglin Zhang, Howard Nusbaum

## Abstract

Intuition and deliberation are two strategies for problem-solving and decision-making. It is commonly believed that deliberation requires more effort than intuition. However, to date, neural evidence approving or disapproving this point is scarce. To explore this issue, we asked participants to play two well-matched games requiring either deliberation or intuition. Using event-related brain potential (ERP) technique, we found: (1) Deliberation elicited more deflective ERP than intuition during −1000 ~ −500 ms before the response, which is consistent with the common belief. (2) More importantly, intuition evoked more deflective ERP than deliberation during 550 ~ 650 ms after the onset of the stimulus, which suggests that intuition may need more effort than deliberation at early stage, contradicting the common belief.

## 1. Introduction

Intuition and deliberative reasoning are two common strategies for human decision-making. Whether individuals make a decision by rational reasoning or intuition is based on different individual characteristics and situations. It has been suggested that there are sustained differences between the individual preference for intuition and deliberation (Betsch and Iannello 2010; Betsch and Kunz 2008). Some people tend to make decisions based on deliberation, while others have a preference for intuition. In addition, people also apply different strategies in different situations (Pachur and Spaar 2015). For example, deliberation is used a lot when playing chess. When two are playing, they must use specific mathematical rules, mature reasoning and deliberative logical thinking to make a rational strategy (Atherton et al. 2003). In contrast to chess in which deliberative reasoning is required to make a decision, there are various other games in which people just need to trust their gut feelings. The famous game “stone, paper, scissors” is a typical example. In this game, individuals guess what action their opponents will take based on their intuitions.

The distinction between intuition and deliberation has become a popular topic in the past few decades. Deliberation is characterized as slow, effortful, controlled, explicit, and rule-governed (Kahneman 2003; 2011), while intuition is “the ability to understand something instinctively, without the need for conscious reasoning” (Oxford Advanced Learner's Dictionary). The distinction between these two strategies or decision-making processes has sparkled much discussion, not only phenomenologically, but also theoretically. Posner and Snyder (1975) distinguished the two processes depending on whether they operated in a controlled or automatic manner. Epstein (1994) contrasted one process, which is experimental and holistic, with the other, which is rational and analytic. Sloman (1996) mapped the two processes onto two distinct cognitive systems: one is associative, whereas the other is rule-based.

One of the essential features that apparently distinguish the two processes is effort. One process is effortful, whereas the other is effortless. Intuition is an emotional judgment or a pure and immediate apprehension (Hayashi 2001; Osbeck 2001). A core property of intuition is that it occurs instantaneously, and thus requires less cognitive effort. The intuitive responses are determined by the accessibility of mental contents. A central concept of intuition analysis is that some particular thoughts come to mind more easily than others (Kahneman 2003). However, deliberation is considered to be an effortful thought that demands much work. Deliberation is associated with higher order control, long-term delay of gratification, and very high levels of abstract thinking (Epstein 1994).

However, is deliberation more effortful than intuition in neural meaning? We consulted many studies in previous literature, but found few neural study that focused on this issue. A functional magnetic resonance imaging (fMRI) study revealed different brain areas were activated when people used deliberation or intuition. The anterior cingulate cortex and the insula showed greater activation when people made an intuitive decision, whereas the middle frontal gyrus and the precuneus were more activated when people solved problem through deliberative reasoning (Kuo et al. 2009). In our view, these results imply that intuition can be more (less) effortful than deliberation in some (other) brain areas, though this point was not explicitly expressed in that paper. In the current study, we asked a further question: can intuition be more (less) effortful than deliberation in some (other) time windows of neural processes?

To explore this issue, we designed two kinds of number games (following and developing the paradigm of (Kuo et al. 2009)), namely, dominance-solvable game and coordination game. The two kinds of games have a similar appearance but a different essence: coordination games require intuition, whereas dominance-solvable games call for deliberation (for more detailed description of these two games, see Methods). A high-density (64 channels) event-related potential (ERP) recording was employed to record the time course of brain electrical activity when people played dominance-solvable or coordination game. More “effortful” means more cognitive resources are required and more neural activities are engaged. According to the popular belief that deliberation is more effortful than intuition, it was expected that deliberation would lead to more deflective ERPs than intuition in entire processes.

## 2. Materials and Methods

### 2.1. Participants

Twenty-one healthy undergraduates participated in this study for payment. All participants were right-handed and had normal or corrected-to-normal vision. They signed written informed consent to participate. This study was approved by an institutional review board at our university. After data collection, we did a careful data check for quality control, and excluded five subjects from analysis: The 1st subject was actually a pilot subject using an old version of procedure; in 3 subjects, the impedances of EEG recording were far out of range (>50 kΩ) or the scalp potentials were unbelievably high (>80 μv) in most trials at many electrodes; one subject did not complete the whole experiment. Therefore, the analysis was based on data of sixteen participants (mean age: 22.0 years; age range: 20 to 23 years; seven females).

### 2.2. Stimuli and procedure

The experiment involved two types of games, which are illustrated in Fig. 1.

**Figure 1.**
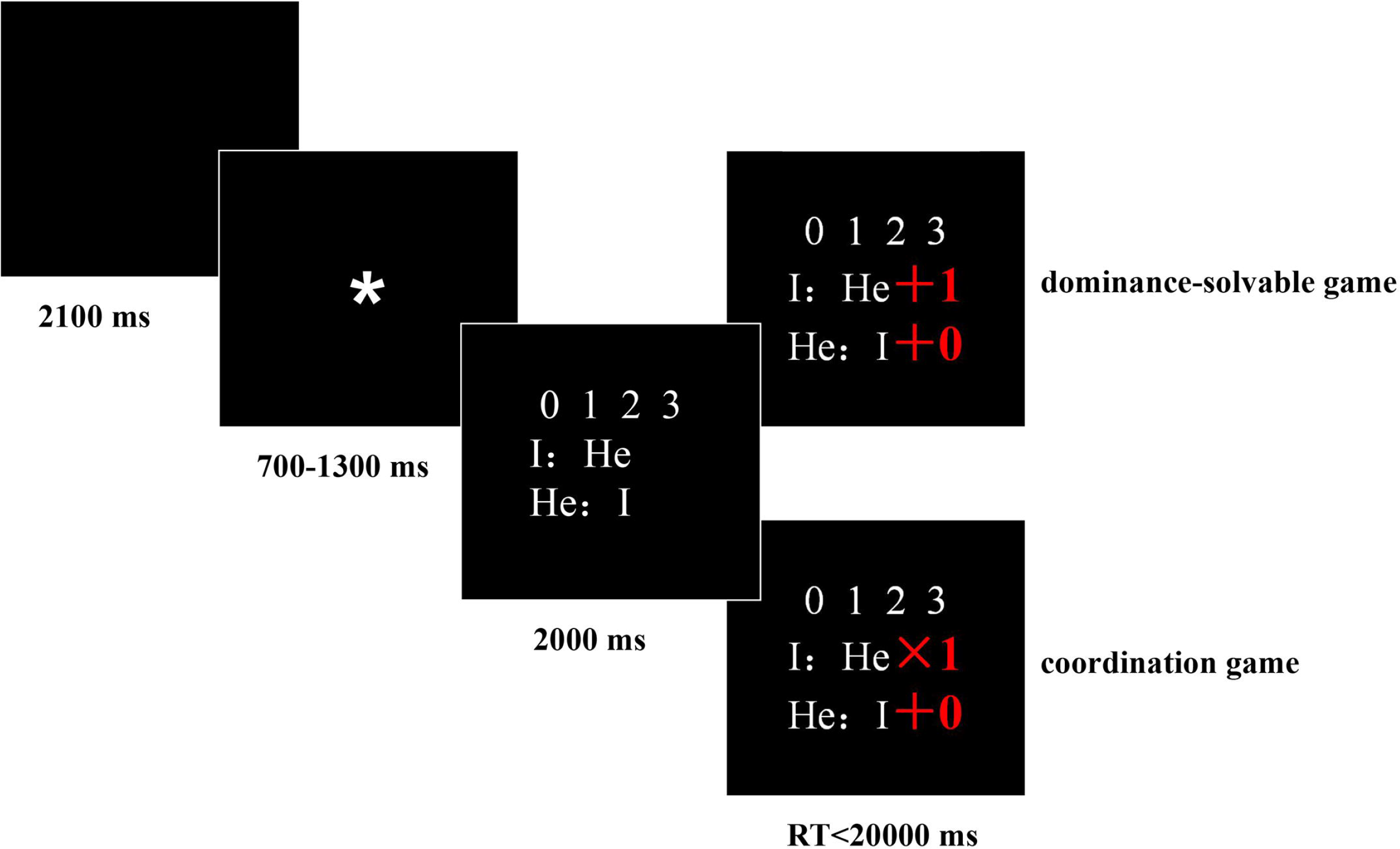
Samples of the two kinds of games. For the two games, the screens were completely identical in the first 3 steps. They were very similar in appearance in the final step, which, however, could incur largely different cognitive processes. See the main text for details.

In the dominance-solvable game, the participants initially saw a black screen for 2100 ms; then, an asterisk (*) appeared in the center of the screen for 700 to 1300 ms to alert the participants that the game was about to begin. Next, four digits arranged in a line and “I: He” with “He: I”, which were arranged vertically, were presented on the screen for 2000 ms. In the fourth screen, the participants could see the targets of “I” (the ERP participant, denoted as A for short hereafter) and “He” (the other player, denoted as B for short hereafter). As shown above, the experimental procedure was divided into four phases and the appearance of the two games was identical for the first three phases, which meant that the participants used the same perceptual and reasoning processes in the first three phases and that the differences only occurred in the final phase. We only analyzed the final phase in which the involvement of the neural processes is different between these two games. As exemplified on the fourth screen of dominance-solvable game, the direction “I: He+1” means A’s (the ERP participant’s) target was choosing a number from 0,1,2,3 that equaled the number chosen by B (the other player) plus 1. The direction “He: I+0” means B’s target was choosing a number from 0,1,2,3 that equaled the number chosen by A plus 0. If a participant or player considered that his target number (the number that was most likely to achieve his target) was not one of the four numbers, he should choose one of the four numbers that was closest to the target number. A could not communicate with B. It was instructed that, if a participant or player achieved his target, he would receive an additional monetary reward. To receive more rewards, A would try his best to reach the goals. First, because A’s target was to choose a number with a value 1 greater than B’s number, A would eliminate 0 (whatever B chose among the four numbers, there was no possibility that 0 was the number that equaled B plus 1). Then, A should surmise B’s choice based on his target. Considering that B’s target was matching A’s choice, A would eliminate 0 for B. Next, A should exclude another number based on his target and the number that he thought B would not choose. Because A’s target was the number 1 greater than B’s number, if B would not choose 0, he would exclude 1. At this point, A has eliminated two numbers (0,1) after three steps. Then, to reach the goal, A only needs to repeat some of these steps. A would not choose 1, and neither would B, whose target was to choose a number equal to A’s. In the fifth step, A realized the best choice for B was 2 or 3. Thus, to choose a number that is 1 greater than B’s number, A would eliminate 2. Thus, A eliminated 0, 1, and 2 and chose 3 as the best number through a six-step reasoning process, which engaged several neural processes, including attention, working memory and mental arithmetic.

In the coordination game, the participant saw the same content as in the dominance-solvable game except on the fourth screen (see Fig. 1). On the fourth screen of coordination game, as an example, the participant saw the following directions: “I: He×1” and “He: I+0”. In essence, A’s target was to choose a number that was the same as B’s, and B’s target was to choose a number that was the same as A’s. Therefore, to obtain more rewards, the participant simply needed to guess the number that the other player would choose.

Questions such as “I: He+1” and “He: I+0” for A were adapted into questions such as “I: He+0” and “He: I+1” for B to match A’s and B’s questions. To explore the ERP difference between the two types of games, either A’s or B’s ERP could be recorded. We focused on A’s ERP. Accordingly, we recorded both the behavioral and ERP data for each A but only the behavioral data for each B. Each A was instructed that a B was playing with him at the same time, although the B players actually played the games at different times. The reason we did not let the two players play the game simultaneously is that we wanted to simplify the programming process on the premise that it would not cause any impact on our experimental results. The number of participants mentioned in the Participants section refers to the number of A’s. The data analysis was based on data of A’s except where data of B’s are explicitly mentioned.

The participant was instructed to choose one of the numbers quickly and accurately by pressing one of four buttons on the keyboard when the targets appeared on the screen. Four right fingers, including the forefinger, middle finger, ring finger, and little finger, corresponded to the four buttons on the keyboard. The participant was told that if he could not choose a number in 20 seconds, the trial would automatically pass to the next one. Thus, the participant lost the chance to get the monetary reward on the trial.

At the beginning of the experiment, each participant was given detailed instructions and examples about how to play the two kinds of games. The experiment comprised 160 trials in which 80 trials were dominance-solvable games and 80 were coordination games. The sequence of the trials was randomized for each participant. To familiarize the participant with the rules of our games and the button pressing before the formal recording, 30 of the 160 trials were used as practice at the beginning of the experiment, which left 130 trials for the formal experiment. The participant took a rest after finishing half of the entire experiment. He was seated in a quiet room where a screen was placed 80 cm away from his eyes. The largest visual angle of the stimuli was 4.8° (horizontal) × 4.0° (vertical). He was instructed to stay still and avoid blinking, except when the totally black screen appeared.

### 2.3. EEG recording and analysis

An elastic cap with electrodes for 64 scalp sites was used to record each subject’s brain electrical activity (Brain Product, Munchen, Germany), with the reference placed on the left mastoid. The vertical electrooculogram (VEOG) generated from the blinks and the vertical eye movements was also recorded using miniature electrodes placed approximately 1 cm above and below each subject’s right eye. The horizontal electrooculogram (HEOG) was recorded using electrodes placed at the right side of the right eye and the left side of the left eye. All electrode impedances were maintained below 10 kΩ. The EEG, VEOG, and HEOG signals were amplified and digitized with a sampling rate of 500 Hz and a bandpass of 0.1–100 Hz.

The EEGs underwent the following steps of offline preprocessing. They were re-referenced to an averaged mastoid reference. The eye movement artifacts (eye blinks and movements) were corrected. The EEGs were filtered with a high cutoff of 16 Hz, 12dB/oct and then segmented and baseline-corrected. The segments whose peak voltages exceeded ±80 μV after correction were excluded before averaging. All of these steps were performed using the Brain Vision Analyzer software (Brain Products).

Our focuses were two windows of time in the decision phase: (1) The early difference between the two conditions. The ERP waveforms were time-locked at the onset of the stimulus in the fourth screen of Fig. 1. The averaged epochs for the ERP were 850 ms, including a 650-ms post-stimulus waveform and a 200-ms pre-stimulus baseline. On the basis of the grand averaged ERP and topographical maps, for the time window of 550 ~ 650 ms, nine electrode points were chosen for statistical analysis: C1, C2, Cz, CP1, CP2, CPz, P1, P2, and Pz.

(2) The late difference between the two conditions. The ERP waveforms were time-locked at the offset of the stimulus (i.e., the time point of response) in the fourth screen of Fig. 1. The averaged epochs for the ERP were 2500 ms, including a 1000-ms pre-offset waveform and a 1500-ms post-offset waveform, with the baseline defined as the post-offset waveform from 1000 to 1500 ms. On the basis of the grand averaged ERP and topographical maps, for the time window of −1000 ~ −500 ms, 20 electrode sites were chosen for statistical analysis: Fpz, Fp2, AF3, AF4, Fz, F1, F2, F3, FCz, FC1, FC2, FC3, Cz, C1, C2, C3, CPz, CP1, CP2, and CP3.

The mean amplitudes were then measured accordingly. For all statistical analyses, the data were computed using repeated-measures ANOVA with the Greenhouse–Geisser correction applied to the p-values.

## 3. Results

### 3.1. Behavioral performance

The mean response time was significantly longer in dominance-solvable games (mean ± sd = 9305 ± 737 ms) than in coordination games (mean ± sd = 3364 ± 524 ms) [*t*(15) = 9.638, *p* < .001], which is consistent with the objective requirement for the dominance-solvable game that engages more neural processes. In the dominance-solvable game, according to the rules of formal game theory, there is an optimal choice for participants. The mean proportion of choosing these optimal numbers was 0.3861, which was significantly higher than the random level (0.25) [*t*(15) = 3.954, *p* < .01]. In consideration of the high difficulty of the game, this result suggested that participants performed the experiment in a serious manner. Furthermore, the mean proportion of the optimal choices in dominance-solvable games showed a significant negative correlation with the steps needed to achieve the target (*r* = −0.512, *p* < .001), which showed that the increment of difficulty is accompanied by the decrease of the “correct rate”.

For the coordination game, technically, there is no way to evaluate whether a choice is correct or not. However, following (Kuo et al. 2009), it is reasonable to use an index, defined as the probability that two randomly chosen participants made the same choice, to evaluate participant success. The formula is as follows:

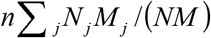

where n means the task has n choices; N (resp. M) means Population A (resp. B) consists of N (resp. M) players; and *N*_*j*_(resp. *M*_*j*_) means *N*_*j*_(resp. *M*_*j*_) players make choice j, for j = 1, 2, …, n.

NCI = 1 means that the players only make a random choice. NCI>1 means that the players’ gut feelings work well. We calculated NCI for each question and then calculated the mean NCI across 80 questions to be 1.0998, which was significantly higher than 1 [*t*(79) = 4.320, *p* < .001]. This outcome suggests that our participants used their intuition to make a choice rather than random guessing.

### 3.2. ERP results

There was an early difference in the decision phase between the two conditions. The ERP waveforms elicited by the two different games at Pz, Cz and CPz are shown in Fig. 2A. A more positive ERP deflection was evoked by the coordination game than by the dominance-solvable game in the 550 ~ 650 ms time window according to the grand average of the ERP and the topographical maps (see Fig. 2A). A 2 (game: dominance-solvable game vs. coordination) × 9 (electrode) repeated-measures ANOVA was conducted (p-values were adjusted using the Greenhouse–Geisser correction). The results revealed the main effect of the game type [*F*(1, 15) = 14.558, *p* = .002, *η*_*p*_^2^ = .493] in the time window of 550 ~ 650 ms. The main effect of the electrode site was marginally significant [*F*(8, 120) = 2.461, *p* = .071, *η*_*p*_^2^ = .141]. The interaction between the game type and the electrode site was not significant [*F*(8, 120) = .687, *p* = .584, *η*_*p*_^2^ = .044].

**Figure 2.**
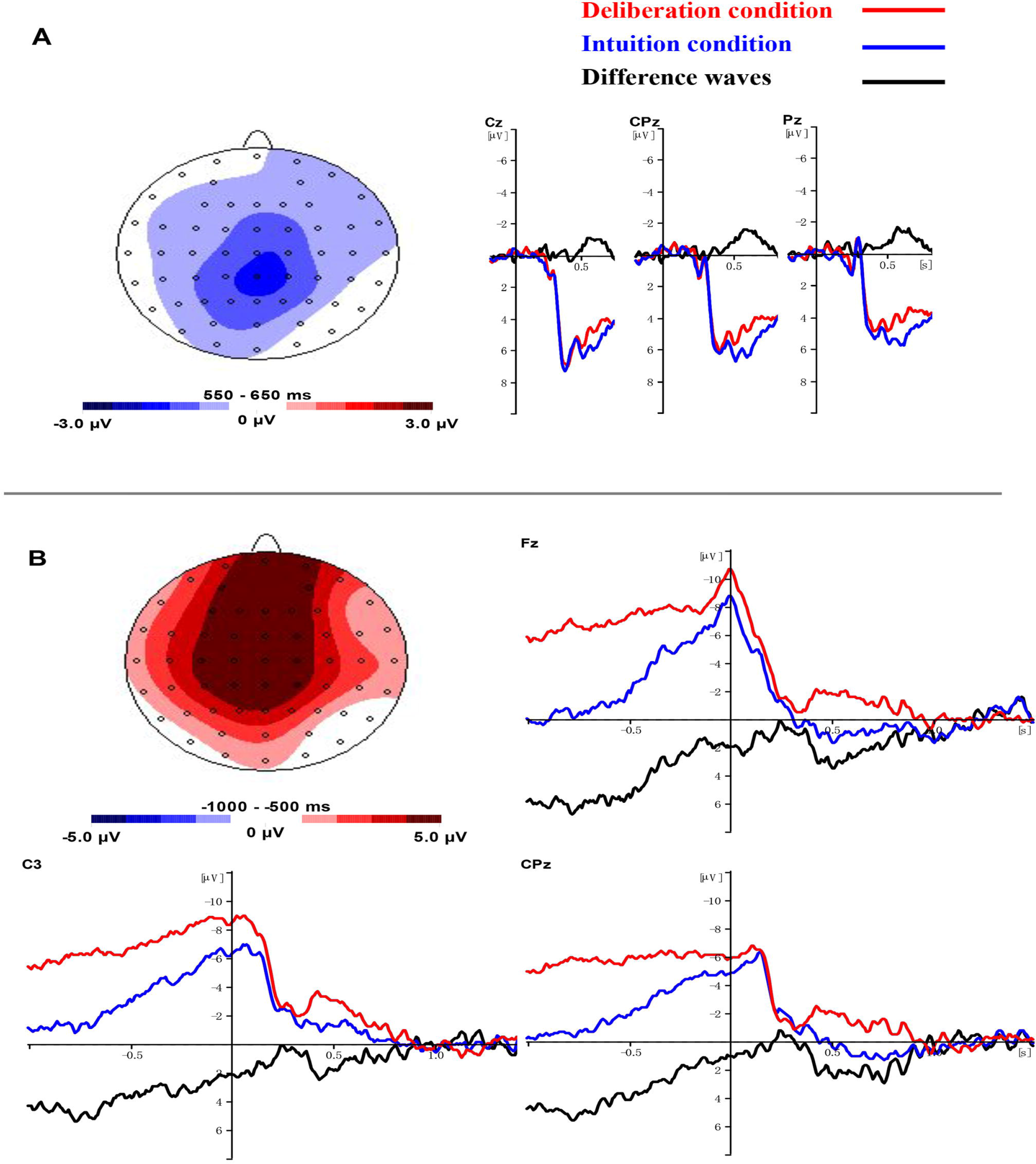
The ERP results. (A) at early stage (550-650 ms after onset of the final screen of Fig. 1), intuition induced more deflective brain potential, as shown by the topography (left) and waveforms (right). (B) at late stage (500-1000 ms before the participants’ response / offest of the final screen of Fig. 1), deliberation induced more deflective brain potential, as shown by the topography (top left) and waveforms (other parts).

There was a late difference in the decision phase between the two conditions (response locked analysis). The ERP waveforms elicited by the two different games at Fz, C3 and CPz are shown in Fig. 2B. A more negative ERP deflection was evoked by the dominance-solvable game within −1000 ~ −500 ms (relative to the time point of response), as shown by the grand average of the ERP and topographical maps (see Fig. 2B). A 2 (game: dominance-solvable game vs. coordination game) × 20 (electrode) repeated-measures ANOVA was conducted (p-values were adjusted using the Greenhouse–Geisser correction). The results revealed the main effect of the game type [*F*(1, 15) = 16.597, *p* = .001, *η*_*p*_^2^ = .525]. The main effect of the electrode site was not significant [*F*(19, 285) = 1.147, *p* = .342, *η*_*p*_^2^ = .071]. The interaction between the game type and electrode site was not significant [*F*(19, 285) = .514, *p* = .710, *η*_*p*_^2^ = .033].

## 4 Discussion

Two major findings emerge from this study: (1) Deliberation elicited a more negative wave than intuition during the −1000 ~ −500 ms before the response, which is consistent with the common expectation that deliberation that engages more neural processes and cognitive demands would lead to higher scalp electrical potentials. (2) Intuition evoked a more positive wave than deliberation during the 550 ~ 650 ms after the stimulus appeared, which violated the common expectation. We will discuss possible reasons for this surprising result in the following section.

Deliberation is described as a slow and effortful process that can provide an analytic and rational decision through step-by-step reasoning. One of the reasons why deliberation is a slow and effortful process is that deliberative process is reflective (Evans 2010). In the dominance-solvable game, to ensure that every strategy they made was optimal, participants needed to exert effort to monitor every step. They adjusted their strategies through this kind of reflective process. More specifically, in the dominance-solvable game, participants first needed to exclude the obviously impossible choice by mental arithmetic. Then, they would surmise the possible choices of the other side by holding the targets of both participants in working memory. Eventually, through these complex and recursive cognitive operations, participants reached an optimal choice. During the deliberative process, because participants needed to make greater and greater efforts to remember the targets, the possible choices and the numbers that have been excluded, the demands on their working memory would increase over time. The growing working memory demands might be one of the reasons for the higher scalp electrical potentials. The evidence from the behavioral results and topographical maps supports this explanation. First, our behavioral result is consistent with previous studies, which showed that the rate of correct responses decreased with the increasing demands on working memory (Gibbs and D’Esposito 2005; Studer et al. 2010). In addition, from the topographical map (Fig. 2B), it is clear that the more negative wave elicited by the dominance-solvable game is concentrated at the front and parietal parts of the brain scalp, which are two regions that are strongly identified with working memory functioning (Courtney et al. 1998; D’Ardenne et al. 2012; Jonides et al. 1998; Narayanan et al. 2005).

Another possible cause of the more negative wave elicited by deliberation is the involvement of mental arithmetic. During the dominance-solvable game, the participants needed to use mental arithmetic to eliminate the impossible numbers. Previous neuroimaging studies have emphasized the critical role of the left parietal lobe in mental arithmetic for number processing (Dehaene et al. 2003; Fulbright et al. 2000). Thus, the increased requirements for mental arithmetic might have evoked the more negative wave at the front and parietal parts of the brain scalp.

The second finding that surprised us was that intuition evoked a more positive deflection than deliberation during the 550 ~ 650 ms after the stimulus appeared. Intuition occurs almost instantaneously (Eisenhardt 1989; Perlow et al. 2002), and does not require effortful step-by-step reasoning which is supported by working memory. Intuitive process merely provides a default solution to a complex problem, like finding a shortcut (Evans 1989; Evans 2006; Stanovich 2010; 1999). However, our result suggests that finding a superior “shortcut” may be an effortful process. First, according to the latency and topographical map (Fig. 2A), we infer that this more positive wave is P3b. P3b is a positive-going ERP deflection peaking at approximately 300 ms (Picton 1992; Pritchard 1981), although its latency is sensitive to the task-processing demand and may vary from 250 to 500 ms (Polich 2007). Therefore, we can infer that the latency of P3b delayed approximately 250 ms in our relatively difficult tasks. Then, the topographical map (Fig. 2A) of our experiment showed that the more positive wave evoked by intuition is concentrated at the parietal part of the brain scalp, which is consistent with previous studies that indicated that the amplitudes of the P3b are classically highest on the parietal brain areas (Polich 2007). A resource allocation hypothesis suggested that the amplitude of this late centroparietal positivity is related to the cognitive resources engaged during the dual-task performance (Isreal et al. 1980; Kok 2001; Wickens et al. 1983). In the dual-task paradigm, when two tasks are performed simultaneously, as the primary task difficulty increased, the amplitude of P3b evoked by the secondary task would decrease, and it would be expected that the extent of this reduction reflects the extent of the cognitive demands associated with the primary task (Isreal et al. 1980; Kramer et al. 1985). Thus, we can infer that the amplitude of P3b can reflect the cognitive demands of the task.

In our experiment, during the 550 ~ 650 ms time window, intuition evoked a more positive P3b than deliberation; thus, it could be presumed that intuition required more cognitive resources than deliberation during this time period. As we know, when they have limited time, instead of engaging in complex reasoning, people always choose to trust their gut feeling. Moreover, a study using rats has suggested that increasing the rats’ sampling time did not result in an improvement in accuracy (Zariwala et al. 2013), which means that even if the decision time is long enough, animals still will not make a decision based on deliberation. Intuition process occurred in a very short time (Eisenhardt 1989; Perlow et al. 2002) but required extracting a lot of information from the previous experiences. This rapid extraction process might play an important role in the reasons why intuition evoked greater P3b in the early phase because the amplitude of P3b is related to the memory-retrieval process (Guo et al. 2006; McEvoy et al. 2001; Rugg and Doyle 1992).

In conclusion, the current study tested the popular belief that deliberation is more effortful than intuition. Our results suggest that both intuition and deliberation are effortful, but they have different effort-consumption dynamics: Deliberation evoked a more deflective ERP than intuition in the late stage (−1000 ~ −500 ms before response), which is consistent with the popular belief; intuition evoked a more deflective ERP than deliberation at the early stage (550 ~ 650 ms after stimulus onset), which suggests that intuition may need more effort than deliberation at early stage, challenging the popular belief.

